# Extreme weather risk shrinks range size estimates and alters biodiversity predictions

**DOI:** 10.1101/2025.10.13.682097

**Authors:** J.M. Cohen, D. Ellis-Soto, F.A. La Sorte, S. Sharma, W Jetz

## Abstract

Extreme weather events, including heat waves, cold snaps, and droughts, are increasing in frequency and intensity with expected but little understood consequences for biodiversity. Extreme weather events can push organisms beyond their physiological thermal or hydric tolerances and thus limit where they can persist, affecting their geographic distributions. Species might be especially sensitive to extreme weather at the edges of their geographic ranges, where they are often already living near their physiological limits. However, the influence of climatic variability and extreme weather is often ignored in favor of climatic means when estimating distributional and richness patterns. Here we link hundreds of millions of citizen science bird observations from 2004-2024 to high-resolution extreme weather risk maps to explore how climatic variability and extreme weather risk alters summer and winter distributions and biodiversity patterns for 535 North American species. We find that species distribution models accounting for historical extreme weather risk performed better at predicting richness and the presence of individual species across 220 well-surveyed sites. Models incorporating extreme weather predicted narrower geographic distributions than models relying on only climatic means, with species’ ranges shrinking an average of 6% in summer and 10% in winter and range truncation observed at the range edges. These effects were observed in both seasons but were particularly strong in winter, a time with greater short-term weather variability than summer. Richness estimates were substantially lower when extreme weather was accounted for, especially in the US southwest and central plains (up to 30-40 fewer species), regions highly prone to extreme heat, cold and drought. Our results suggest that more mechanistically informed biodiversity predictions that account for extreme weather are critical for reliably predicting shifting distributional and biodiversity patterns.

## Introduction

### Extreme weather: an underappreciated driver of species distributions and biodiversity patterns

Anthropogenic climate change has been linked to rapid changes in ecological patterns and processes, including species re-distributions (Pecl et al., 2017) and phenological shifts (Cohen et al., 2018), and is threatening one out of six species with extinction (Urban, 2015). While most work to date has explored the ecological consequences of gradually rising mean temperatures, the influence of increasingly frequent and intense extreme weather (Chen et al., 2025; Coumou and Rahmstorf, 2012; Masson-Delmotte et al., 2021) for wildlife species and biodiversity patterns has been largely underexplored (Astigarraga et al., 2024; Gonçalves et al., 2024; Soifer et al., 2025). While organisms are typically adapted to handle predictable seasonal or daily variation in environmental conditions within their ranges (Vázquez et al., 2017), extreme weather events occur unpredictably with anomalous conditions at a far greater magnitude than typical environmental variability (Cattiaux and Ribes, 2018). Many organisms are physiologically constrained by their upper and lower thermal limits (Buckley and Huey, 2016) and thus may be more likely to be pushed past their physiological limits by an extreme weather event than gradually shifting mean conditions over decades or expected variability (Germain and Lutz, 2020; Harris et al., 2018; Strangas et al., 2019). Thus, understanding how extreme weather events limit the boundaries of species distributions and niches is critical for accurate ecological forecasts and better conservation outcomes under rapidly accelerating climate change (Ellis-Soto et al., 2023; Maxwell et al., 2019; Parmesan et al., 2000; Van De Pol et al., 2017).

Previous works exploring how organisms respond to extreme weather highlight substantial species-specific responses to extreme heat and drought. Species with generalist diets and increased mobility tolerate extreme heat better than sedentary species with specialized diets (Astigarraga et al., 2024; Cohen et al., 2021, 2020). Waterbirds appear more resilient to extreme heat and cold events during their migratory periods, with the pressure to arrive at their breeding grounds possibly outweighing the costs of moving to avoid extreme weather events (Masto et al., 2022). On the contrary, raptors such as Golden Eagles (*Aquila chrysaetos*) equipped with telemetry devices appear to be highly sensitive to both periods of extended drought and heat, altering their daily behavior and space use with negative consequences for their survival (Wiens et al., 2018). Endotherms, such as birds and mammals, may reduce their activity time for foraging and reproduction to avoid extreme thermal conditions exceeding their physiological tolerances (Huey et al., 2012; Riddell et al., 2019). The combination of periods of extended drought and heat can lead to increased thermal and hydric stress with detrimental effects for wildlife (Fuller et al., 2014; Prugh et al., 2018). With the increasing availability of remote sensing reanalysis products with high spatial and temporal resolution (Anderson, 2018; Chen et al., 2024; La Sorte et al., 2021), it is now possible to incorporate extreme weather risk when estimating species distribution and biodiversity patterns.

### The importance of range edges

As species are typically near the limits of their thermal tolerances at the edges of their distributions (Fredston et al., 2021), extreme weather in these regions may push species past their physiological limits and influence their ability to survive and reproduce (Soifer et al., 2025). Thus, estimated range limits may shrink when accounting for extreme weather while range interiors remain relatively intact (Maclean and Early, 2023). This may be especially true in deserts and continental interiors where weather variability is high and heat waves, droughts and cold snaps are more common, although organisms in these regions may have adaptations to tolerate or avoid extreme conditions. Alternatively, range edges can also serve as critical refugia under certain extreme conditions (Soifer et al., 2025). For mobile taxa, range edges can serve as critical refugia during climate extremes: for example, Dickcissels (*Spiza americana*) irrupt into northern edge habitats when drought at the core-range ‘pushes’ them northward (Bateman et al., 2015). Therefore, range edges could plausibly see reductions or expansions under extreme weather.

### Aims

Here, we leveraged observational data from eBird, a global citizen science platform in which users submit bird sightings, to explore how species’ distributions and biodiversity patterns are modified by the inclusion of variables describing extreme weather risk. We utilize species distribution models (SDMs), which relate environmental factors to species occurrence information and are commonly used to understand the abiotic factors limiting the range of a species as well as distributional and biodiversity patterns in space (Guisan and Thuiller, 2005). However, traditional SDM approaches relating occurrence information to climatic means (Elith and Leathwick, 2009; Lawlor et al., 2024) cannot differentiate between habitats that differ in weather variability - for example, a desert that experiences both cold and warm extremes and a coastal region that experiences constant intermediate temperatures. While SDMs sometimes include metrics of weather variability including diurnal temperature range or seasonality, these metrics might fail to describe the risk of a species eclipsing its thermal tolerance (sensu Taheri et al., 2024), especially if the predictable daily or seasonal variability falls within the thermal limits of the species (Briscoe et al., 2023).

We modeled the seasonal distributions of 535 North American bird species in both summer and winter to explore how extreme weather risk influences predictions of species distributions and continental-scale patterns of biodiversity during multiple times of year. We fit SDMs including only mean climate variables (*Means* models) and compared them against models containing means and climate variability (*Variability* models) and means, variability, and extreme weather risk (*Extremes* models). To represent extreme weather risk, we leveraged two recently developed high-resolution data products: global, seasonal extreme heat and cold risk (La Sorte et al., 2021) and monthly drought risk (Chen et al., 2024). Models lacking variability or extreme weather information included simulated data with similar autocorrelation structure to ensure any improvement in prediction was not artifactual. We evaluated predictions across all three model types using both model cross-validation and by comparing richness, presence, and absence predictions to observations at 220 well-sampled, evenly distributed sites around the continent. Finally, we explored how the importance of extreme weather to distributions is moderated by species’ phylogenetic relationships and functional or life history traits, including body size, hand-wing index and habitat preference.

We expected that estimated range sizes would shrink for most species with the inclusion of information about climatic variability and extreme weather risk in models and that most change in model prediction would occur around the range edges. We expected that if richness predictions differ between *Means* and *Extremes* models when distributional predictions are combined, this would primarily occur in areas with high risk of extreme heat, cold, and drought such as the Great Plains and American Southwest (Diffenbaugh et al., 2008). We expected any effects to be strongest during winter, when many species are already living at the edge of their thermal tolerances and have few options to fulfill dietary, habitat and niche requirements following a disturbance (Casson et al., 2019; Penczykowski et al., 2017). Finally, we expected that small-bodied species, those with a small hand-wing index or those not from forest habitats might have distributions that are more sensitive to extreme weather as they are less likely to have the thermal inertia, movement capabilities, or habitat buffering required to persist under such conditions, respectively (Huey et al., 2012; Jarzyna et al., 2016).

## Methods

### Modeling area and species selection

We modeled the distributions of bird species native to the United States and Canada within a region between 179.99° to 42.68°W longitude and 0° to 55°N latitude. We excluded northern Canada and Alaska because spatial data coverage is poor and predictions can be unreliable. It was necessary to model into Central and South America to generate accurate predictions for species with only a small portion of their ranges in the southern US.

We compiled a list of 689 native bird species that annually breed or overwinter in the United States and Canada based on the most recent version of AviList (Rheindt et al., 2025). We included American Birding Association birding codes (Pyle et al., 2024) that distinguish regularly occurring species (code 1 or 2) from vagrants (code 3+). Some species from our original list were excluded because their ranges within our study extent (mainland US and Canada) are almost exclusively marine, are Hawaiian endemics or had an insufficient number of records (<50 post-thinning; see below) for analysis. Species were only modeled for one season if they do not occur in the US or Canada in the other season. In the end, we modeled 535 unique species, modeling summer distributions of 481 species and winter distributions of 486 species.

### Data acquisition, filtering, and thinning

In July 2024, we compiled observational data from eBird, a global citizen science initiative in which users submit checklists containing bird observations. eBird data has become widely used for understanding species distributions (Sullivan et al., 2014). Users have the option to indicate whether all observed species were recorded on the checklist, marking checklists as “complete”, which allows the inference of absence points and presence-absence modeling. Although absence points represent detection rates and not true absences, they are typically expected to be consistent within species if accounting for time of year and time of day, and so uncertainty in detection should not bias the current study. Users also indicate the level of effort involved in each observation by providing the distance traveled, amount of time birdwatching and number of observers (hereafter, effort indicators), each of which is accounted for in our models (see below). We separately compiled all ‘complete checklists’ submitted to eBird that were recorded during the summer (June-August) and winter (December-February) for almost all species. For shorebirds (Charadriiformes), we limited data to June so their long-distance July movement would not affect distribution predictions, which are meant to approximate breeding ranges. For any other species with a mean August latitude of presence observations > 2° latitude (*ca*. 222 km) lower than the June-July mean, indicating significant migration in August, we excluded August observations and used only June-July observations.

In all checklists, we discarded subspecies information and applied several filters to the data to reduce bias and improve data quality following established eBird data modeling protocols (Johnston et al., 2021; Kelling et al., 2018). We first eliminated checklists covering extreme high survey durations (> 3 hours), with high numbers of observers (>5 individuals) or adhering to any protocol other than “stationary” or “traveling”, as these comprise only a small subset of the data (< 0.5%) and are difficult to compare with the bulk of the dataset. Second, we reduced the positional error of observations (Gábor et al., 2020) by eliminating checklists covering > 2 km because these are likely to result in high spatial uncertainty, mismatching the true observation location with environmental coordinates. Third, we removed data before 2004 as there is too little data from earlier years to adequately control for long-term temporal population trends. We limited overprediction outside of the species’ range boundary by limiting checklists to those falling within a 200 km buffer of the species’ seasonal expert range boundary (via Cornell’s spatial boundaries - https://ebird.org/science/status-and-trends/download-data, accessed May 2024; Ebird, 2024).

Citizen science datasets typically suffer from spatial and temporal biases in reporting, as users typically report observations in populated areas and during popular times of year for birdwatching (Steen et al., 2019). To reduce site-selection and temporal bias in data collection and better even out the spatial and temporal distribution of the data, we thinned observations to one per 5 km grid cell per week, as recommended by eBird’s best practices guidelines (Johnston et al., 2021; Strimas-Mackey et al., 2020). We repeated this thinning for checklists in which the species does and does not occur to reduce imbalance between presence and absence points (Robinson et al., 2018). Species were required to have at least 50 observations post-thinning to be included in the analyses. To address uneven observer effort between checklists, we controlled for duration of survey event, distance traveled, and number of observers in all models (see below). Such filtering and thinning decisions do not impact the centroid or spatial autocorrelation of the data (which could influence how presence observations are associated with environmental conditions), only partially evening out data density in space and time (Cohen and Jetz, 2025).

All coordinates, polygons and grids used in the study operated under a conical equal area projection. The *raster* (Hijmans et al., 2015) and *sf* (Pebesma, 2018) R packages were used for spatial geoprocessing.

### Model covariates

We included several classes of covariates in the SDMs to account for a variety of factors that could influence species distributions. First, we included temporal covariates – year, date, and time of day – in all models to account for temporal variability in bird activity and long-term population trends, and we included all effort indicators as covariates to address uneven sampling effort between checklists. Topographic, landcover, and climate covariates were selected from a pool of 32 contenders based on minimal collinearity. All covariates were spatially aggregated to 1 km based on the mean value in each cell. All topographic and landcover covariates were included in all models, while the climatic covariates included depended on the model category (*Means, Variability,* and *Extremes*).

Our topographic suite of covariates included mean elevation (from EarthEnv; Robinson et al., 2014), seasonal mean enhanced vegetation index (EVI; MODIS; https://lpdaac.usgs.gov/products/mod11a1v006/), and terrain roughness index (TRI; EarthEnv). Our landcover suite of covariates included percent landcover type based on mean-aggregating all 300m cells falling within the 1km cell. Landcover information corresponded to the year the checklist was recorded for the following categories: cropland, evergreen broadleaf, deciduous broadleaf, evergreen needleleaf, deciduous needleleaf, mixed forest, mosaic, shrubland, grassland, lichens/mosses, sparse, flooded/freshwater, flooded/saltwater, flooded/shrub, urban, barren, water, and ice (ESA CCI, 2017).

Climatic covariates were selectively included based on model type so we could compare models including only mean climate variables, climate variability metrics, or those describing extreme weather risk. *Means* models included only mean annual temperature (bio1) and mean annual precipitation (bio12) from CHELSA v2.1 (Karger et al., 2021). *Variability* models included those two variables and temperature seasonality (bio4) and precipitation seasonality (bio15) from CHELSA. Finally, *Extremes* models included the above variables plus seasonal extreme heat event (EHE) and extreme cold event (ECE) duration risk via (La Sorte et al., 2021) based on the number of consecutive days containing EHE or ECE during each season. EHE and ECE maps were derived from the European Centre for Medium-Range Weather Forecasts (ECMWF) fifth generation atmospheric reanalysis of the global climate (ERA5) database (see La Sorte et al., 2021 for details). We used the maximum duration of EHE and ECE for each season, year, and grid cell averaged across years. *Extremes* models also included mean seasonal Standardized Precipitation Evapotranspiration Index (SPEI; Chen et al., 2024) to estimate drought risk. SPEI is available as a historic monthly product at 1 km resolution, so to create mean seasonal versions we averaged monthly SPEI for either June-August (summer) or December-February (winter) from 2004-2018 (the available years that span the eBird data we used) to generate seasonal average SPEI across North America. *Means* and *Variability* models additionally included simulated layers with similar autocorrelation structure to corresponding variability/extreme weather layers absent from those models (see below).

### Prediction surface

We generated a prediction surface covering our bounding box at 1 km spatial resolution containing values for all the environmental covariates (topographic, landcover, and climate) corresponding to each cell. To limit prediction of distributions to the approximate region where each species occurs, we masked portions of the prediction surface that fell outside of the buffered range extent when predicting the distribution of each species. We generated predictions for non-spatial covariates assuming a 1 km, 1 hour search with 1 observer for the year 2023 (the final year with 12 months of data) at a randomized date within the season and during the hour of the day when the species is detected most often.

### Species distribution models

Our goal was to explore how climatic variability and extreme weather alter species distributions. We modeled the distributions of each species using Random Forest, a machine learning method designed to analyze large datasets with many covariates which is often found to generate the most accurate SDMs (e.g., Mi et al., 2017). Random forest (RF) is a widely-accepted algorithm to model species distributions using eBird data in recent years (Callaghan et al., 2022; Cohen et al., 2021; Coleman et al., 2018; Johnston et al., 2021) that can flexibly adjust to complex, nonlinear relationships and consider high-order interactions among predictors (Evans et al., 2011). RFs can integrate presence and absence observations and outperform other algorithms (e.g., Maxent) as SDMs (Mi et al., 2017; Zhao et al., 2022) with eBird data (Johnston et al., 2021; Robinson et al., 2018). RFs can sometimes be found to overfit, but this is generally less of a concern when using presence and absence data (Valavi et al., 2021) and when larger numbers of observations are available (Luan et al., 2020), as in the present study (total observations ranged from ∼27,000 to ∼1.3 million depending on species with a median of ∼336,000).

Prior to modeling, we split data 75/25 into training and testing samples using spatial block cross-validation via the blockCV package (Valavi et al., 2018). For each model, 32 equally-sized blocks were generated over the geographic area containing presence observations of each species. Training data was sourced from 24 blocks, testing data from 8, and the identity of each block was systematically assigned such that the proportion of presence to absence points was roughly consistent between the two sets. This was repeated across four folds, with each of the 32 blocks contributing to the testing set in one of the folds, such that all data was in a testing set in one fold and training set in the other three folds. Folds were each required to have at least 10 presence points in each of the training and testing set to run. We further split the training and out-of-bag (OOB) sets randomly 75/25 within each fold. Comparisons of Moran’s I across training and testing sets revealed no differences in spatial autocorrelation (Table S1).

Using the ranger package (Wright et al., 2018), we fit three models – *Means*, *Variability*, and *Extremes* – for each species during both summer and winter seasons, resulting in 2,901 separate models. All models contained all topographic, landcover, and temporal predictor suites as well as effort indicators. *Means* models contained mean annual temperature and precipitation. *Variability* models additionally contained temperature and precipitation seasonality.

Finally, *Extremes* models additionally contained seasonal EHE, ECE and SPEI. Because these additional variables may result in improved performance of *Variability* and *Extremes* models by explaining random error, we simulated dummy layers with a similar autocorrelation structure to each added variable (seasonality and extreme weather) using the *prioritizr* package (Hanson et al., 2025). We included all dummy layers in *Means* models and the extreme dummy layers in *Variability* models.

Models were fit to the training samples and parameterized to 100 trees and 7 threads. Observation points in the training set varied from thousands to millions depending on species. We used the testing set for internal cross-validation, predicting occurrence and non-occurrence based on the covariate values associated with each sample, and compiling model predictive performance metrics including area under the curve (AUC), Kappa, true skill statistic (TSS), sensitivity, and specificity for every fold. For each fold, we also checked test-set calibration plots to diagnose overfitting. We used OOB samples to predict occurrence and non-occurrence for a given species from the occurrence rate, estimating the threshold that maximized the sum of sensitivity and specificity. Following each model fit, we predicted the model to the prediction surface to generate thresholded grid-level occurrence predictions. We further compiled the predicted area of occurrence and variable importance scores for covariates within each fold. We compiled the predictive performance metrics and variable importance scores for each model by averaging across folds. When combining range estimates across species for biodiversity predictions, we used only the thresholded range prediction from the 4^th^ fold.

For all models, we also calculated the proportion of the predicted range that falls within the geographic boundaries of the expert range (generally filled-in polygons delineating the range boundaries) to understand how much overprediction exists beyond the range boundary. While expert range polygons are notably inaccurate for identification of suitable habitat in range interiors, they are generally reliable for approximating range edges. We generated partial dependence plots using the pdp package (Greenwell, 2017) to explore how each species shifted its probability of occurrence relative to each of the three extreme weather risk factors while controlling for all other covariates in the SDM. Plots were generated based on *Extremes* models for each season. To understand how extreme weather constrains the mean climate at which each species occurs, we created plots of mean temperature or precipitation against the corresponding extreme weather risk (EHE or SPEI) for both presence and absence points and 95% best fit ellipses for each.

### Comparing results between model types

We tested if estimated proportion of maximum estimated range area, proportion of prediction within expert range, model performance (AUC and TSS), and contribution of environmental factor type to the model differed among model type (*Means*, *Variability*, *Extremes*) or season using heteroscedastic one-way ANOVAs for trimmed means (Wilcox, 2011) via the *WRS2* R package (Mair and Wilcox, 2020). We used the percentile *t* bootstrap method with a 20% trim level and 10,000 bootstrap samples. We report the percentile *t* bootstrap test statistic as *T* and followed significance tests (p < 0.05) with the corresponding bootstrap post hoc tests.

### Comparisons across species and regions

We fit Phylogenetic Generalized Least Squares (PGLS) models to understand species-level associations between traits and changes to distributional limits while estimating a phylogenetic signal in our results using the *nlme* package (Pinheiro et al., 2017) and an avian phylogeny from (Jetz et al., 2012). We fit four models estimating relationships between traits and the change in either range size or prediction outside of range edges between either *Means* and *Variability* models or *Means* and *Extremes* models.

Traits included as model covariates were trophic level, habitat preference (categorical predictors), log-transformed body mass and hand wing index (continuous) from AVONET (Tobias et al., 2022). Season was additionally included as a two-level covariate. To reduce the number of habitat types and ensure sufficient representation across categories, we combined “forest” and “woodland” habitat categories under “forest” (n=209); “marine”, “coastal”, “riverine”, and “wetland” under “water” (n=155); “grassland”, “rock”, and “shrubland” under “open” (n=155); and reclassified “human modified” to “urban” (n=16). In the trophic level category we condensed “carnivore” and “scavenger” under “carnivore” (n=323) alongside “herbivore” (n=100) and “omnivore” (n=112).

We fit PGLS models using a backward stepwise model selection in which insignificant predictors were iteratively removed until the best fit (according to AIC) was found. We ran the PGLS model on 100 trees drawn from the posterior distribution of the avian phylogeny. Models fit with a Pagel’s λ correlation structure outperformed the simpler Brownian motion assumption because λ estimates the degree of phylogenetic signal rather than assuming it is fixed. Finally, we fit ANOVAs to any final models containing categorical variables to test for significance.

### Biodiversity estimates

We summed either thresholded or relative occurrence rate predictions of seasonal species distributions for all species within every cell in our prediction surface to produce two versions of species richness. We repeated this for each of the three model types. Biodiversity visualizations were generated using color palettes from the *Rcolorbrewer* package (Neuwirth and Neuwirth, 2011).

### Site-level Validations

We additionally validated all predictions by comparing them to observations at 220 roughly evenly-spaced, well-sampled sites throughout the US and Canada. To ensure even spatial representation, we selected the most heavily visited eBird location (minimum 100 visits per season) within every cell of a 200 km grid within our modeling extent (Fig. S1). We first compiled the total number of species and identities of all species observed at every site for each season. To avoid comparisons against inflated richness totals due to occasional vagrancy, species were considered present if observed on at least 2% of checklists and were otherwise labeled as absent. We then compared site richness and species observations against the predictions at each model type and season by calculating R^2^ values. We calculated the percentage of species that models correctly or incorrectly predicted to be present or absent for the 1 km cell corresponding to each observation site (true positives rate and true negatives rate, respectively).

## Results

### Model validations

For both summer and winter seasons, species’ predicted distributions were generally more accurate in models that accounted for climatic variability and extremes. We compared model-based estimates of species presence, absence, and richness to records at 220 well-sampled, evenly-spaced sites around the US and Canada (Fig. S1). We found that *Extremes* models did best at correctly identifying species that were truly present at sites, on average correctly identifying 5% more species per site than *Means* models for both seasons (Fig. 1a,c), which was consistent across the spectrum prediction accuracy (Fig. S3a-d). All models were virtually equally skilled at predicting species absent from sites, averaging correct predictions for 96% of species in summer and 97% in winter (Fig. 1b,d; see Fig. S2 for false presence/absence rates), also consistent across prediction accuracy (Fig. S3e-h). We found that in both seasons, richness predictions based on *Extremes* models best matched observations, with predictions being especially strong for winter (Fig. S4). AUC values based on spatial-block cross-validation were high across model types (typically 0.85-0.95), but we observed a trend towards greater model AUC for *Variability/Extremes* models in winter relative to *Means* models (T = 2.8, p = 0.066; Fig. S5) though not summer (T = 1.4, p = 0.24; Fig. S5).

**Figure 1.**
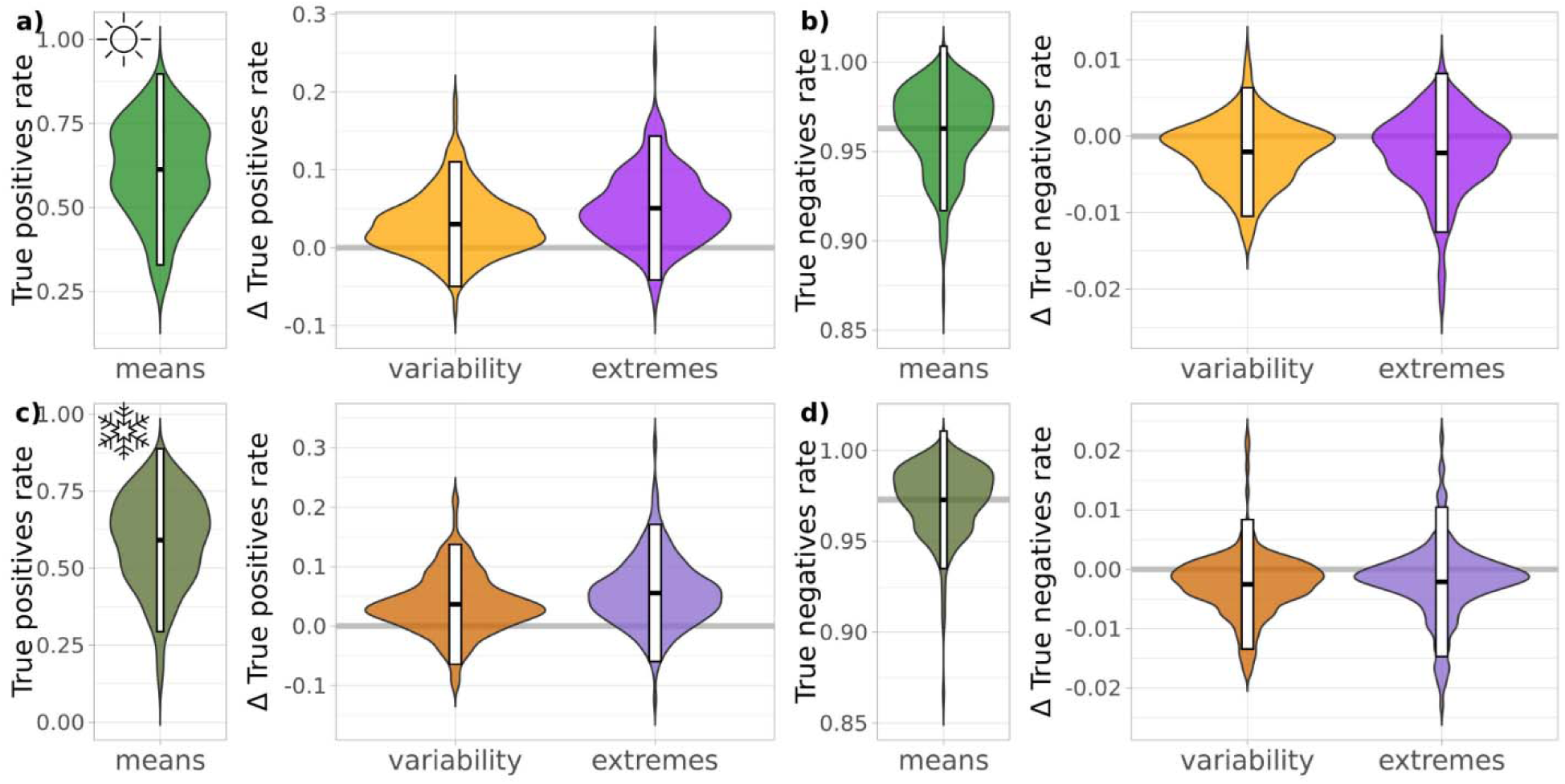
Performance of biodiversity predictions. Violin plots show the proportion of species correctly identified as present or absent at a set of 220 locations across the US and Canada for species distribution models with climate means only (green; a,c, true positive rate; b,d, true negative rate) and the difference in model performance between models fit with climatic variability (orange) and extreme weather (purple) compared with means-only models (see Fig. S8 for false positive/negative rates). Species distributions were modeled for summer (a-b, brighter colors) and winter (c-d, darker colors). White bars give 95% intervals and black lines are medians.

### Range sizes and edges

We found that models including only climatic means overpredicted species’ ranges compared to those accounting for climatic variability and extremes. For example, models including only climate means for hooded oriole (*Icterus calculus*), a species that breeds in the southwestern US, predicted its breeding distribution to be in hot, dry portions of west Texas and interior California that were not supported by models including climate variability and extreme weather information (Fig. 2a for binary occurrence predictions; Fig. S6a for probability of occurrence).

**Figure 2.**
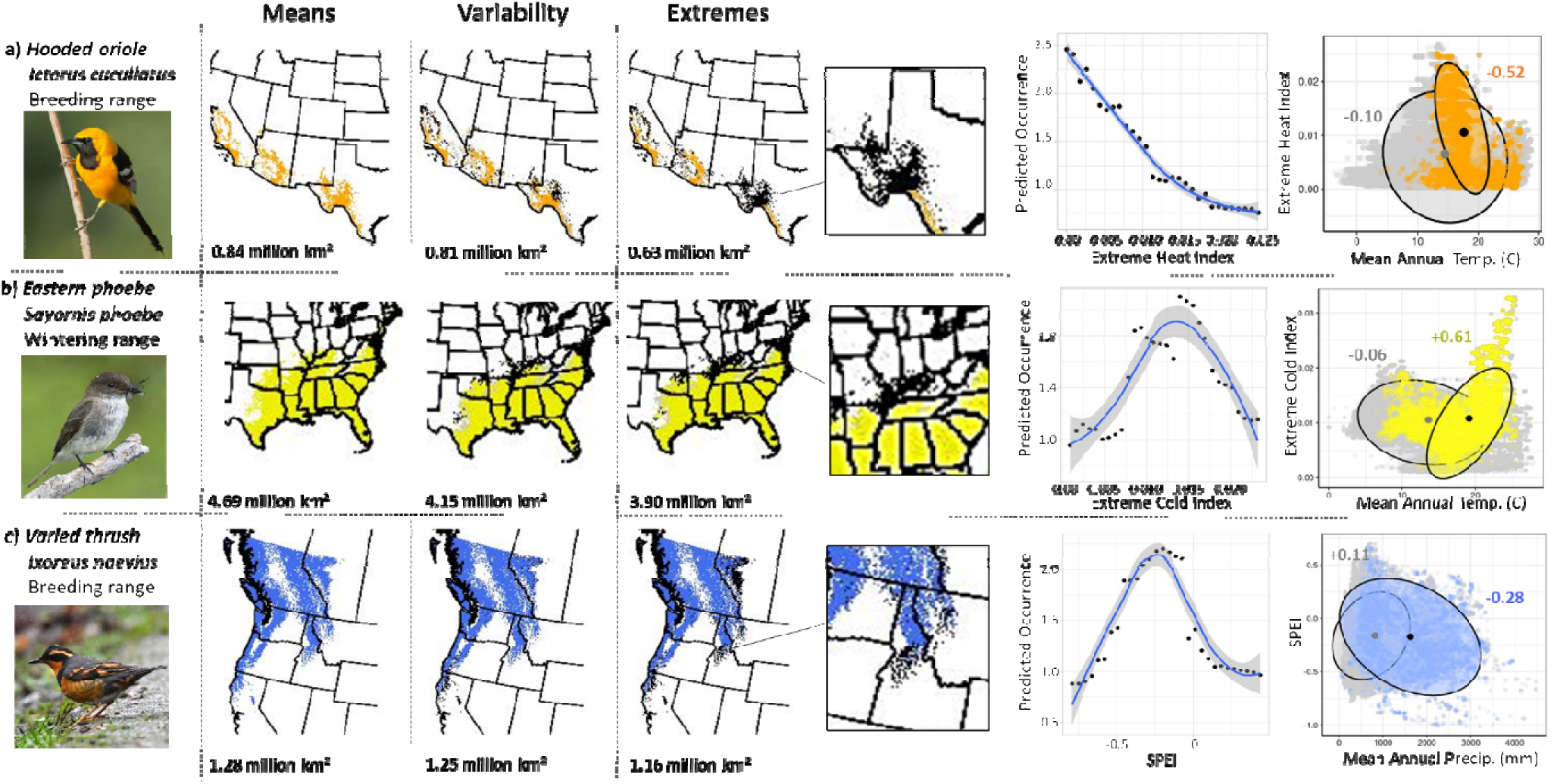
Bird distributions and environmental niches in the context of extreme weather. Predicted breeding range of (a) hooded oriole (*Icterus cucullatus*), (b) wintering range of eastern phoebe (*Sayornis phoebe*), and (c) breeding range of varied thrush (*Ixoreus naevius*) across models including only climate means, climate variability and extreme weather risk (left to right). For variability and extremes, colored areas represent predicted suitable areas in all models, black areas are suitable in *Means* but not *Variability* or *Extremes* models, and pink areas are suitable in *Variability* or *Extremes* but not *Means* models. Insets highlight regions with large differences between model predictions. In the right columns, partial dependence plots convey the estimated relationship between the extreme heat index, extreme cold index, or SPEI and occurrence likelihood. Niche plots (95% ellipsoids) compare a mean climatic condition with a corresponding risk of extreme weather, with colored points and ellipses representing presence observations, gray points and ellipses as absence observations, black points as centroids, and values are linear slopes based on the observations underlying the ellipses, color-coded to presence or absence observations. Mean annual temperature is compared with extreme heat or cold risk and total annual precipitation with drought risk. See Fig. S1 for probability of occurrence estimates for these species.

Including these variables revealed a negative association between extreme heat risk and probability of occurrence for the species in partial dependence plots (PDPs; Fig. 2a). For eastern phoebe (*Sayornis phoebe*), which winters in the southeastern US, PDPs suggested a negative association between species occurrence rates and extreme cold risk and models without extreme weather predicted a larger distribution in northern areas of the map (Fig. 2b; Fig. S6b). Finally, for varied thrush (*Ixoreus naevius*), a species breeding in temperate rainforests of the pacific northwest, PDPs suggested sensitivity to drought risk and models without drought risk overpredicted the range edge to drier regions in California and Idaho (Fig. 2c; Fig. S6c).

Across 481 species, summer range sizes shrank by a median of 5% in *Variability* models and 6% in *Extremes* models across species, and estimates from both model types were marginally different than those from *Means* models (1-way ANOVA: T = 217, p < 0.001; Fig. 3). Winter range sizes significantly shrank by a greater amount, 8% for *Variability* and 10% for *Extremes* (T = 452, p < 0.001). In summer, the proportion of the estimated range falling within the expected boundary (expert range map) increased by 2% in *Variability* models and 3% in *Extremes* models relative to 78% in *Means* models (One-way anova: T = 5.3, p < 0.01; Fig. 4a). In winter, it increased by 4% in *Variability* and 5% in *Extremes* models relative to 70% in *Means* models (T = 5.1, p < 0.01; Fig. 4b).

**Figure 3.**
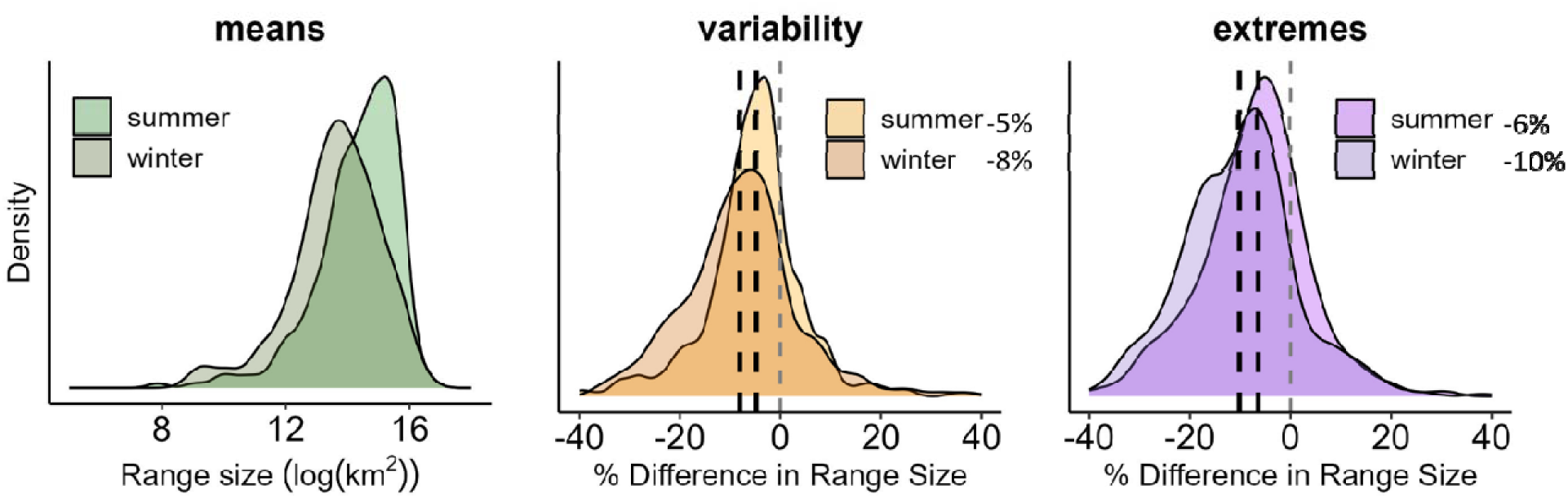
Range sizes are smaller when extreme weather risk is included in models. Density plots show difference in estimated range size between models containing only climate means (green) and those with climate variability (orange) and extremes (purple) including extreme heat and cold event risk and Palmer drought severity index. Summer ranges are represented by brighter colors and winter ranges by darker colors. Black dotted lines are the seasonal medians and gray dotted lines represent zero.

**Figure 4.**
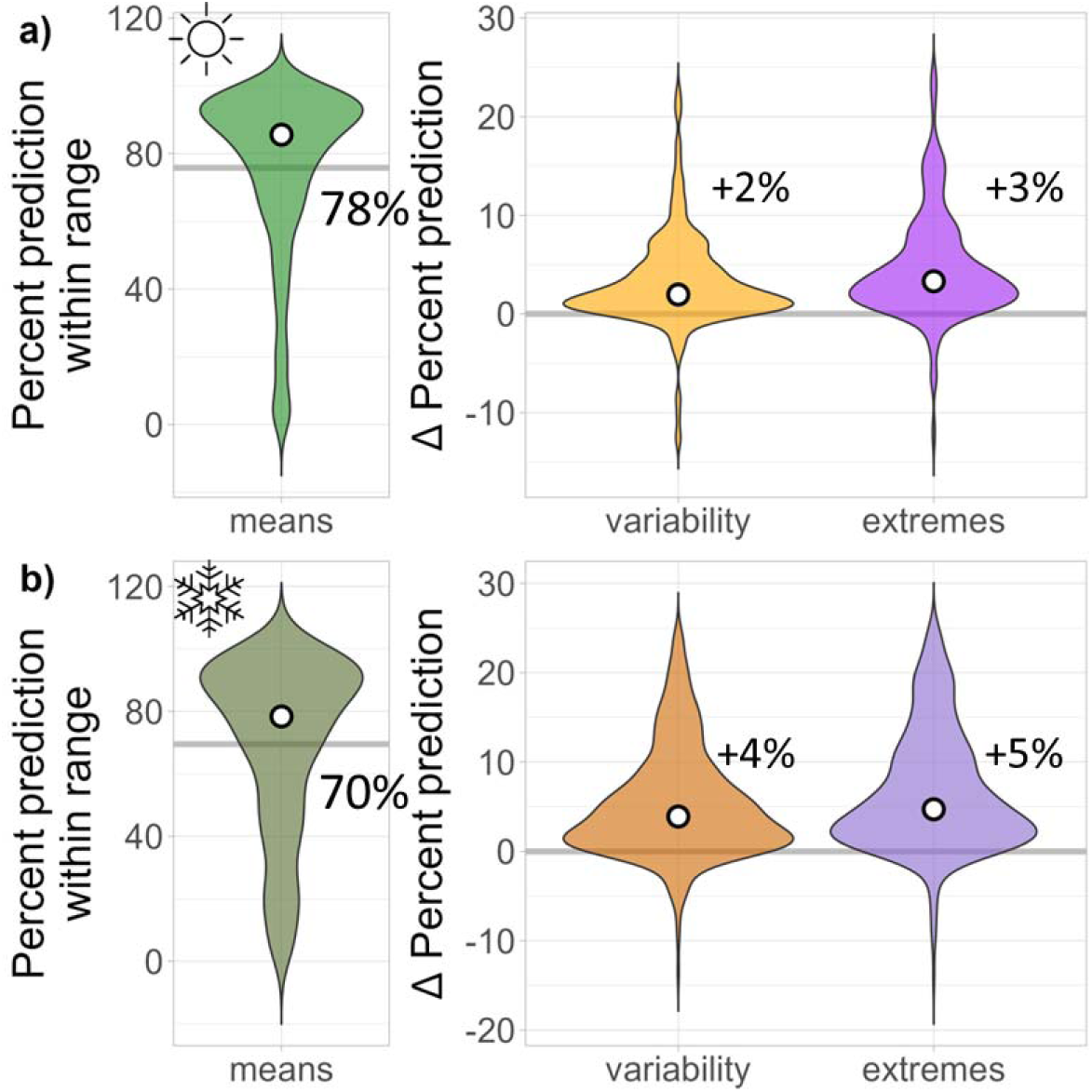
Percent prediction inside the range boundary increases when extreme weather risk is included in models. Violin plots present the change in the proportion of the predicted range that falls within the expert range (a metric of how well overprediction is limited at range edges) between models containing only climate means (green) and those with climate variability (orange) and extremes (purple) including extreme heat and cold event risk and Palmer drought severity index. Higher delta values represent less prediction outside of the expert range, suggesting overprediction. Summer ranges (a) are represented by brighter colors and winter ranges (b) by darker colors. White points represent medians.

The contribution of climate variables to total model predictor importance in Random Forest increased at every step between *Means*, *Variability* and *Extremes* models in summer (T = 46, p < 0.001; Fig. S7a), while the contribution of topographic (T = 26, p < 0.001; Fig. S7b) and landcover variables (T = 29, p < 0.001; Fig. S7c) decreased. We observed the same pattern for winter models as well (climate: T = 66, p < 0.001, Fig. S7d; topographic: T = 34, p < 0.001; Fig. S7e; landcover: T = 60, p < 0.001, Fig. S7c).

### Percent difference from mean models given in figure legends

### Functional Traits and Phylogeny

Our model selection approach using PGLS revealed that body mass was selected as the only final predictor variable across all modeling contexts and was positively associated with changes to range sizes between *Means* and *Extremes* models (β=0.10, p < 0.05), but not *Variability* (β=0.04, p = 0.34; Tables S2-S3). Body mass was also positively associated with changes to the proportion of predicted range falling inside the expert range between *Means* and *Variability* (β=0.11, p < 0.05; Table S4) or *Extremes* (β=0.11, p < 0.01; Table S5) models. Thus, the prediction outside the range margins changed the most between *Means* and *Extremes* models for large bodied species. Across all models, Pagel’s λ correlation structure fell between 0.14 – 0.31, indicating a weak but detectable phylogenetic signal (λ = 0 corresponds to no phylogenetic signal, λ = 1 to a strong signal under Brownian Motion).

### Biodiversity predictions

Richness predictions based on stacked thresholded predictions of species distributions varied between model types. In summer, *Variability* and *Extremes* models expected reduced richness compared with *Means* models primarily in the northern and central Great Plains region (Fig. 5a for delta richness between models; Fig. S8a for absolute richness). In winter, *Variability* and *Extremes* models expected reduced richness in the southwestern US and southern/central Great Plains (Fig. 5b; Fig. S8b). We observed similar trends based on relative occurrence rate estimates (Figs. S9-S10).

**Figure 5.**
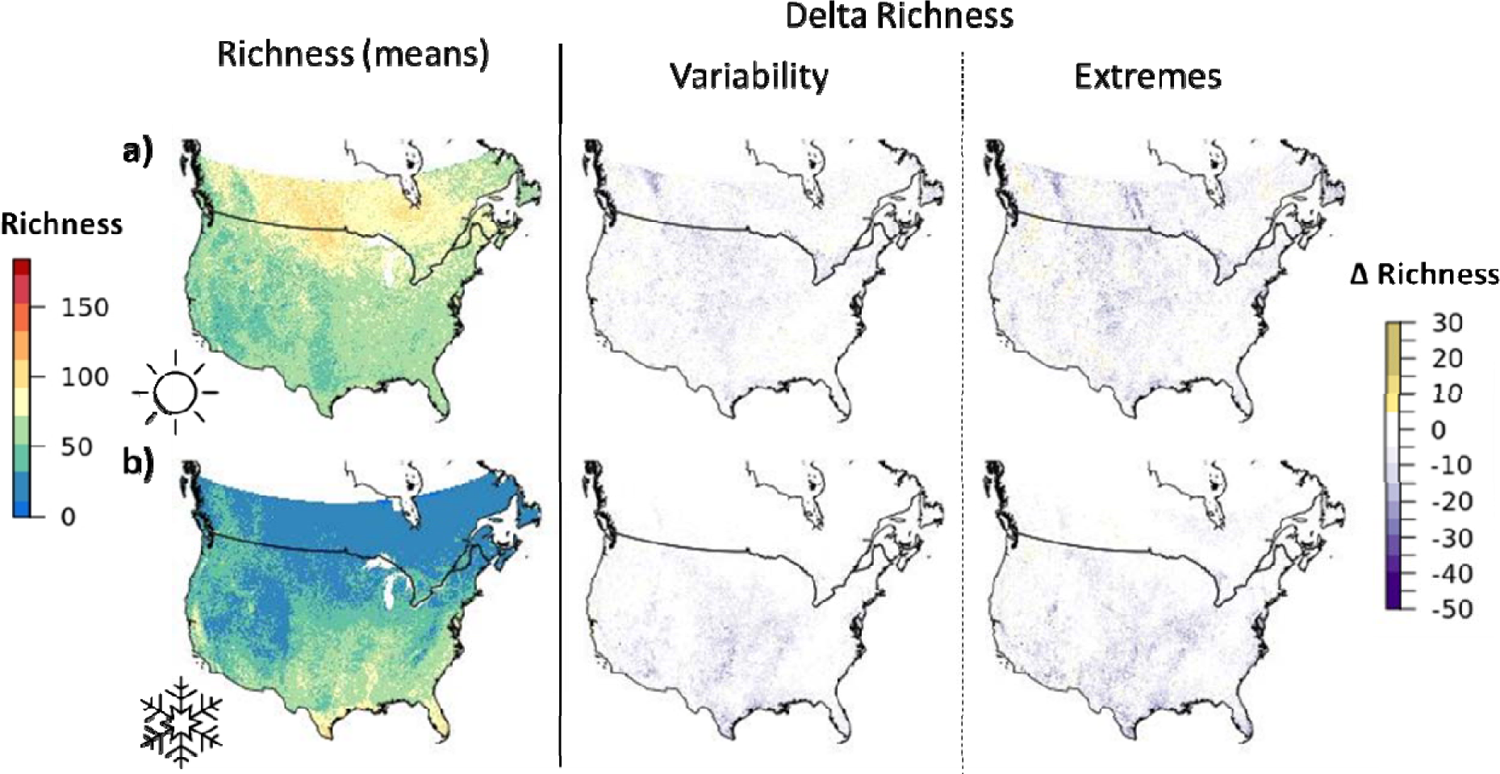
Avian species richness across models containing climate means, variability, and extremes. Left panels show predicted richness based on stacked models containing only climate means for (a) summer and (b) winter. Middle and right panels show the difference between predicted richness based on models containing climate variability and extremes compared with means only, with darker purple areas experiencing a larger reduction in estimated richness and yellow areas experiencing an increase in richness.

## Discussion

Using an integrative species distribution modeling approach for 535 bird species, we show that incorporating extreme weather risk maps improved predictions and consistently altered estimates of species range sizes across both summer and winter. Models considering extreme weather events demonstrated increased performance in cross-validation and when predicting species richness and presence at well-sampled sites (Fig. 1). Range sizes contracted by 5-10% when variability and extremes were incorporated into models compared with those based on climatic means alone (Fig. 2). Truncation of species ranges was larger in winter than in summer (Fig. 3), especially around range edges (Fig. 4) and somewhat surprisingly for large-bodied birds. When assessing the importance of extreme weather on aggregated biodiversity patterns, incorporation of extreme weather events was particularly relevant for the central prairie and southwestern United States, biogeographic regions that are highly prone to extreme weather (Fig. 5).

With an alarming 30% decline in North American bird populations over the past 50 years (Johnston et al., 2025; Rosenberg et al., 2019), it is critical to better understand how increasingly unpredictable and extreme weather affects species’ distributions and biodiversity (Fredston et al., 2025). This study highlights how extreme weather—beyond gradual shifts in mean climate—compresses species’ viable habitats, revealing that traditional methods considering only mean climatic conditions are overestimating distributions by up to 20-40% for some species and biodiversity totals sometimes by dozens of species. Future work could incorporate land-use types alongside future climatic means, variance and extreme-weather metrics (Powers and Jetz, 2019; Zurell et al., 2018) or fine-scale microclimate variation (Maclean and Early, 2023) to yield more mechanistic predictions incorporating multiple drivers of biodiversity change—enabling us to pinpoint which functional traits and habitat configurations will best buffer different bird species against increasingly hot climates.

### Avoiding overprediction of distributions and biodiversity with relevance for conservation

Without accounting for extreme weather, there is a risk of SDMs overpredicting species distribution and patterns of biodiversity, hindering their utility for conservation efforts (sensu EllisuSoto et al., 2021). Incorporating extreme weather metrics may be especially advantageous for predicting species distribution across geographic regions experiencing increasing magnitude and severity of extreme weather. For example, hooded oriole, a heatusensitive species, lost over 10% of their predicted breeding range in interior California and west Texas when extremeuheat risk is included, and eastern phoebe, a southern wintering species that avoids cold climates in winter, saw a 20% contraction of its northern wintering boundary under extremeucold scenarios (Fig. 3). SDMs with extreme weather layers predicted lower estimates of species richness, particularly across ecoregions prone to more frequent droughts, heatwaves and cold snaps, including the American Southwest and Great Plains in the Midwest of the United States. We also found that incorporating extreme weather into models may be most important for specific species depending on their body size. Predicted range sizes were more limited by extreme weather in smaller-bodied birds as expected, as they have relatively low thermal inertia and might be more sensitive to weather fluctuations (Huey et al., 2012), though large-bodied birds surprisingly demonstrated greater predicted range shrinkage around the range edges when extreme weather was included in models.

## Limitations

Several limitations should be considered while interpreting our results. First, the extreme weather risk layers used in our analyses highlight existing areas of high risk but do not identify regions that may see more extreme weather in the future (Seneviratne et al., 2021). Second, these layers do not distinguish the temporal scale of extreme weather events, but sustained events may be most likely to negatively impact organisms (Murali et al., 2023). Third, our models are fit at the species level, but local populations of widespread species may be more or less adapted to handle extreme weather in areas that see more of it, highlighting the need to integrate ensemble approaches that model species-environment relationships locally (Cohen et al., 2023). Finally, our approach does not consider the potential buffering effects of microclimates for avoiding the influence of extreme weather, allowing persistence in regions with high extreme weather risk (Kemppinen et al., 2024), although this may be a more important consideration for taxonomic groups such as mammals, reptiles, amphibians or insects that heavily utilize microclimates (Kerr et al., 2025).

## Conclusions

Recurrent extreme weather–including heat waves, cold snaps, and droughts– may disproportionately truncate range edges and reduce overall habitat suitability. Here, we find that integrating extreme weather risk maps into species distribution models led to model improvements, refined geographic predictions of range sizes, and sizable changes to biodiversity predictions. By explicitly accounting for extremes, our approach yields improved species distribution predictions and richness maps and incorporates one of the major drivers of biodiversity loss into ecological forecasting (Trisos et al., 2020), ultimately, improving species vulnerability assessments to climate change. Moving forward, this framework is readily transferable to ectotherms, whose thermal tolerances and distributions are much closer tied to environmental extremes (Paaijmans et al., 2013). Our findings also highlight the importance of integrating extreme weather into projections of species’ future ranges and suggest that models ignoring these episodic stressors risk painting an overly optimistic picture of species’ persistence (Urban, 2015). Furthermore, incorporating extremeuweather models with highuresolution microclimates could allow researchers to identify thermal refugia —small pockets of habitat buffered against regional extremes—that could serve as conservation priorities under rapid climate change (Hannah et al., 2014). As extreme weather becomes more frequent and intense, our findings pave the way towards a more holistic picture of how such events influence species distributions and biodiversity patterns.

## Supplementary Figures

**Figure S1.**
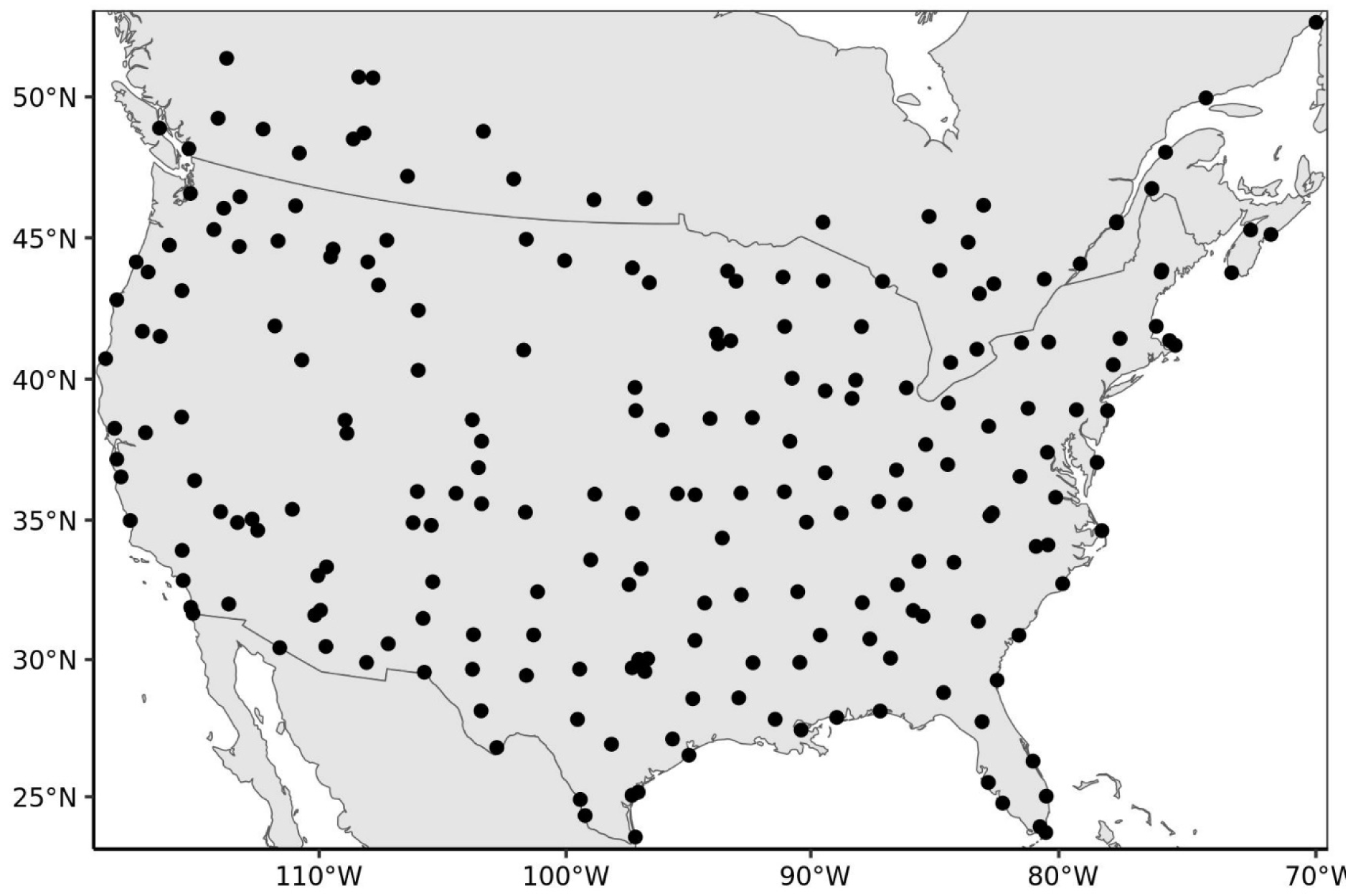
Map of 220 sites used for site-level validations. Sites were selected if they were the most-well sampled site within every 200km grid cell and had a minimum of 100 visits from eBird users

**Figure S2.**
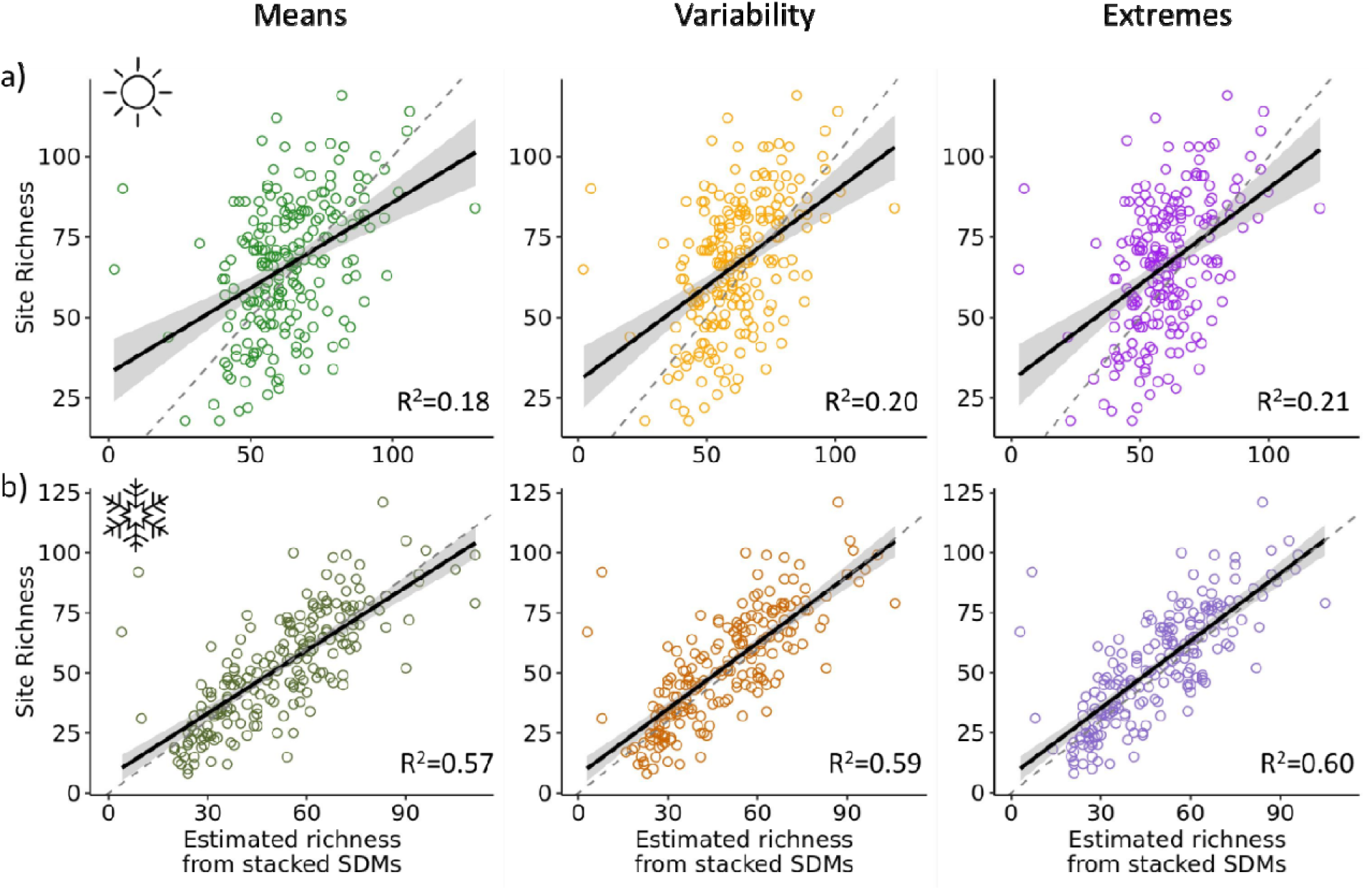
Predicted vs. observed species richness at 220 test sites. We compared the estimated richness totals (x-axes) based on SDMs with only climate means (green) or those including variability (orange) and extreme weather (purple; left to right) against observed richness totals (y-axes) at 220 well-sampled sites across the US and Canada (see Fig. S1 for map) in summer (a, brighter colors) and winter (b, darker colors). The solid line is the line of best fit, gray shading is 95% confidence interval, dashed line is a 1:1 reference line, and associated R^2^ values are provided.

**Figure S3.**
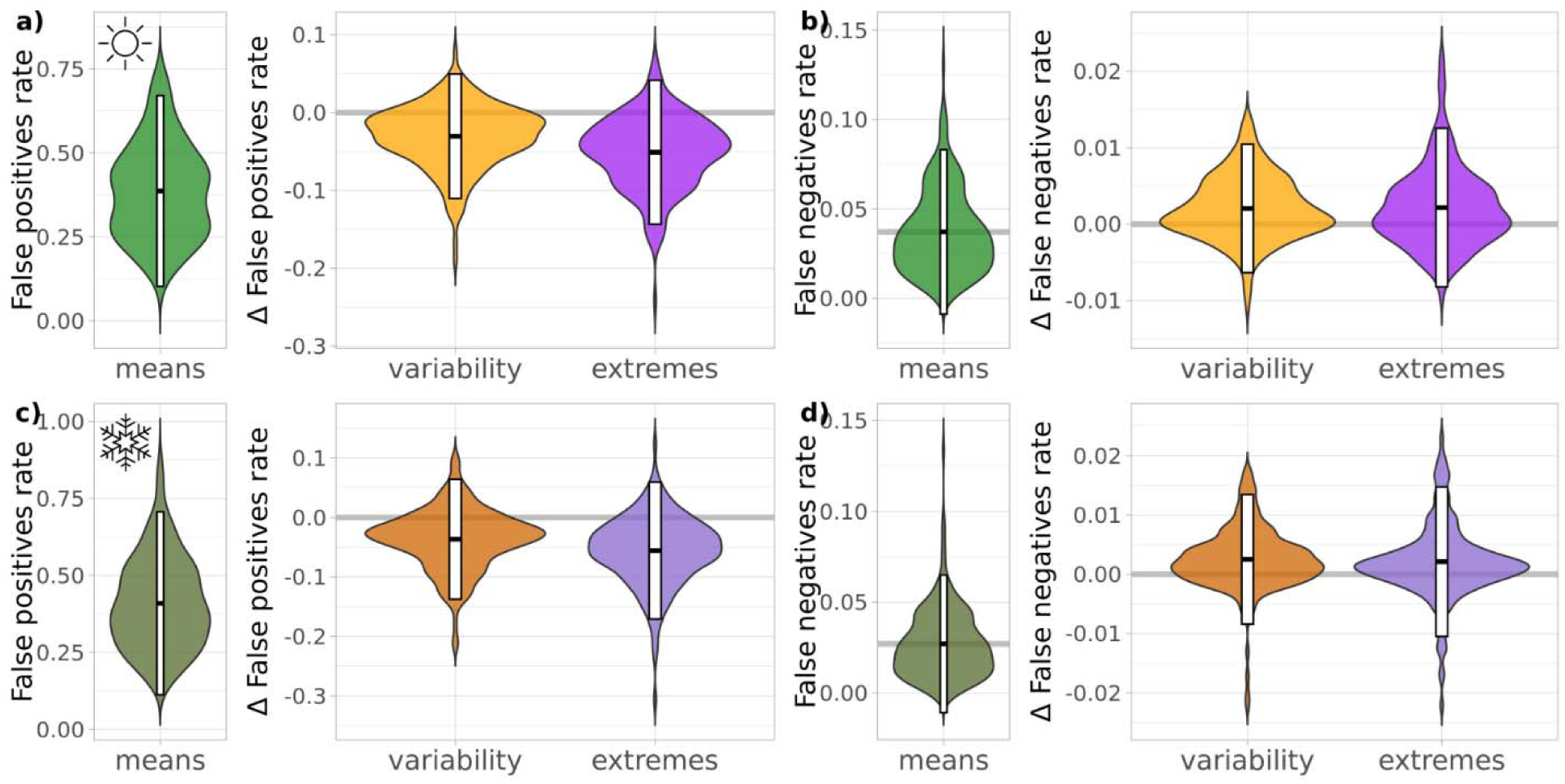
Proportion of species incorrectly identified as present or absent at 220 sites across the US and Canada. Rates are given for models including only climatic mean variables (green; a,c, false positive rate; b,d, false negative rate) and the difference in rates between models fit with climatic variability (orange) and extreme weather risk (purple) included compared with means-only models. Species distributions were validated for summer (a-b, brighter colors) and winter (c-d, darker colors). White bars give 95% intervals and black lines are medians.

**Figure S4.**
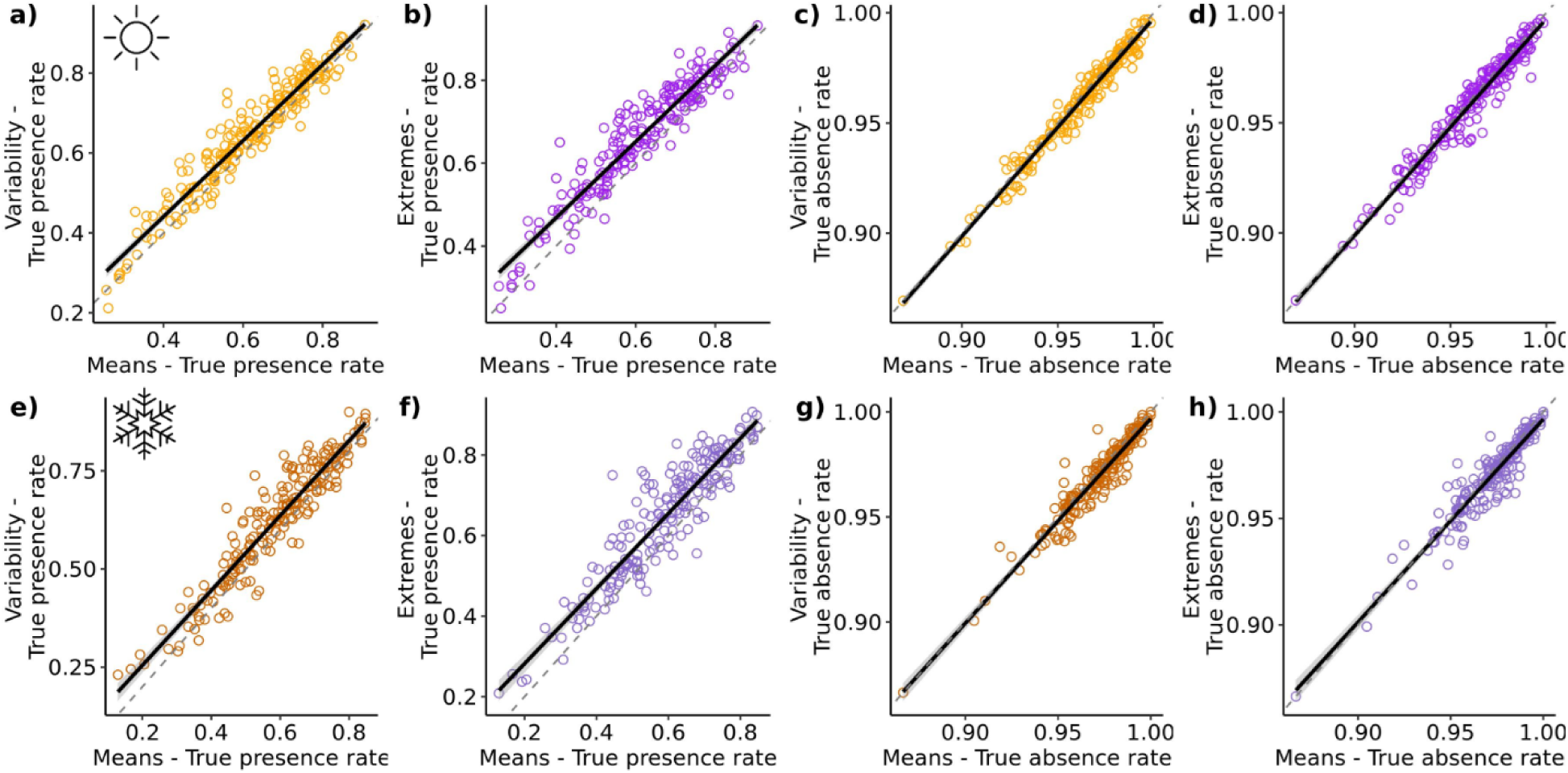
Comparison of true presence and absence rates across model types. For summer (a-d; brighter colors) and winter (e-h; darker colors), true presence rates (a-b,e-f) or absence rates (c-d,g-h) across 220 sites based on models with climate means (x-axes) are compared against those from models with variability (a,c,e,g; orange) and extreme weather (b,d,f,h; purple) on y-axes. Presence or absence rates are calculated by comparing observed and model-predicted species at all sites. Points represent sites, black lines are lines of best fit, gray shading is associated 95% confidence interval, and dotted line is a 1:1 line.

**Figure S5.**
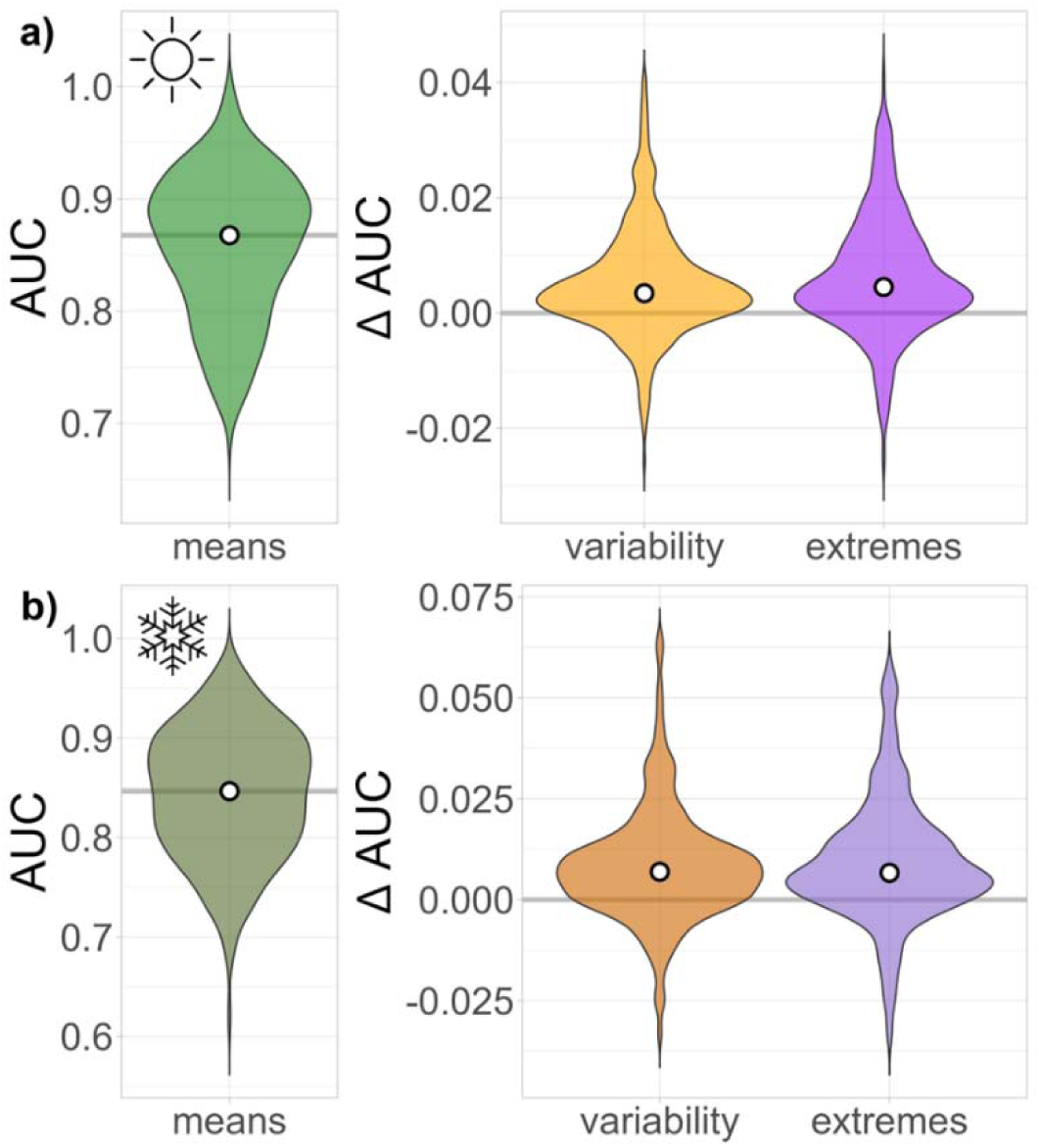
Model performance across models containing climate means, variability, and extremes. Change in AUC between models containing only climate means (green) and those with climate variability (orange) and extreme weather (purple). Summer distributions (a) are represented by brighter colors and winter ranges (b) by darker colors. White points represent medians.

**Figure S6.**
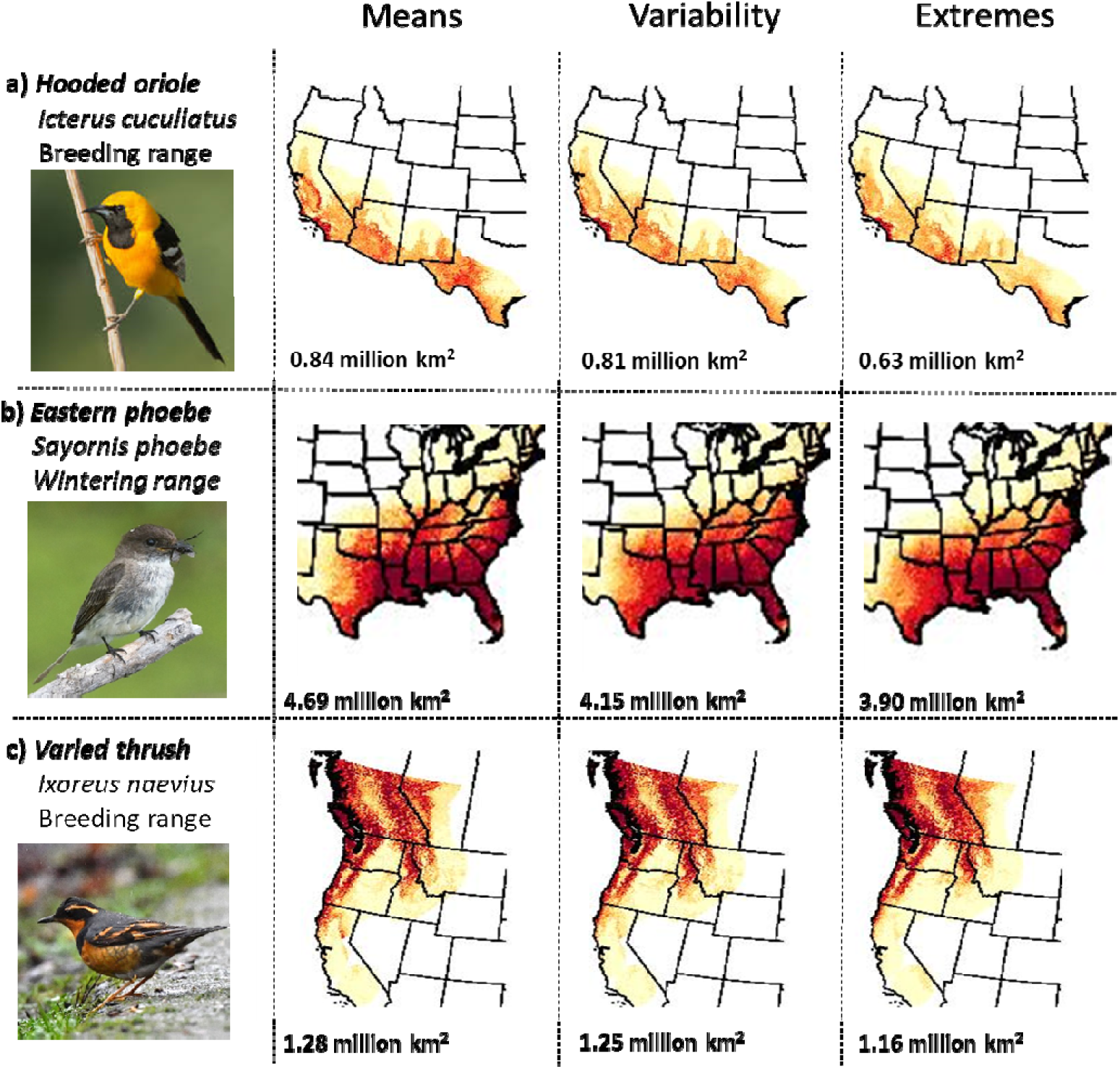
Bird distributions and niches in the context of extreme weather. Predicted probability of occurrence for breeding range of (a) hooded oriole (*Icterus cucullatus*), (b) wintering range of eastern phoebe (*Sayornis phoebe*), and (c) breeding range of varied thrush (*Ixoreus naevius*) across models including only climate means, climate variability and extreme weather risk (left to right). Redder colors signify greater probability of occurrence.

**Figure S7.**
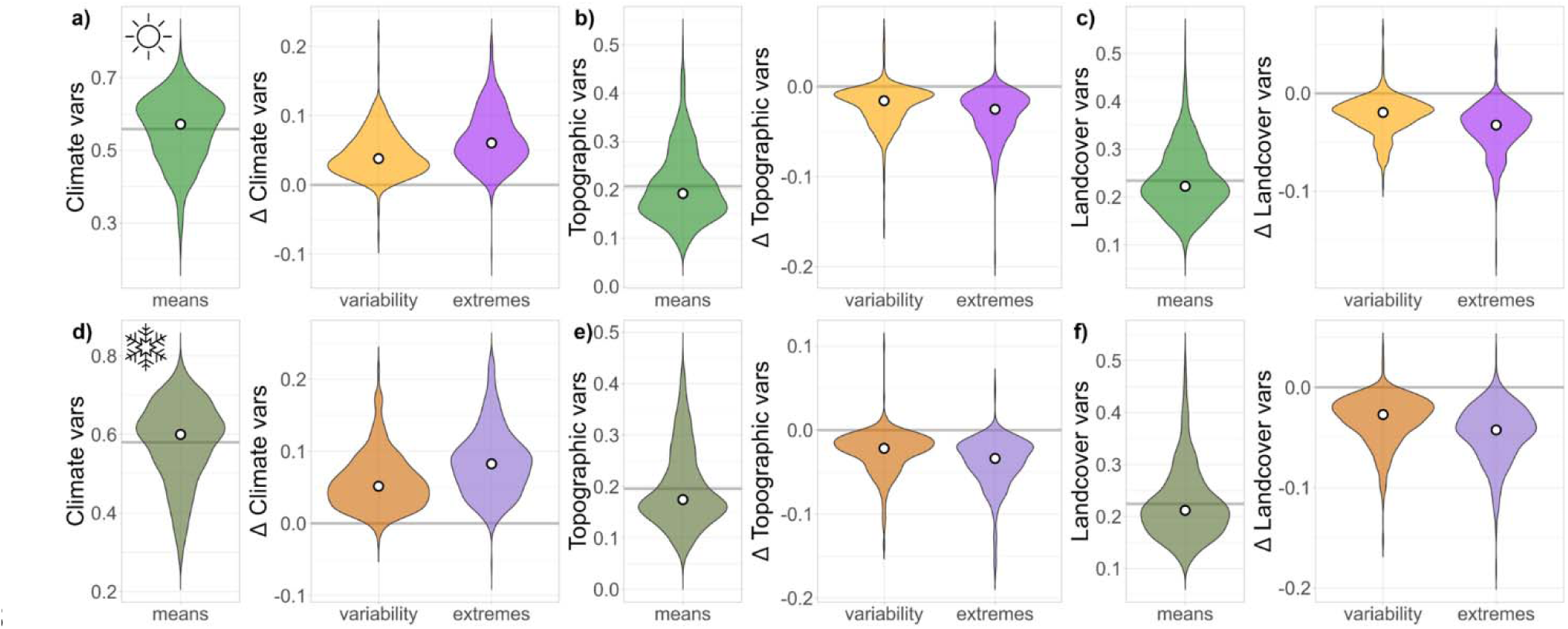
Environmental variable importance across models containing climate means, variability, and extremes. Change in relative contribution of (a,d) climate, (b,e) topographic, or (c,f) landcover variables to the models (based on proportion of total relative importance scores from random forest) between models containing only climate means (green) and those with climate variability (orange) and extreme weather (purple). Summer ranges (a-c) are represented by brighter colors and winter ranges (d-f) by darker colors. White points represent medians.

**Figure S8.**
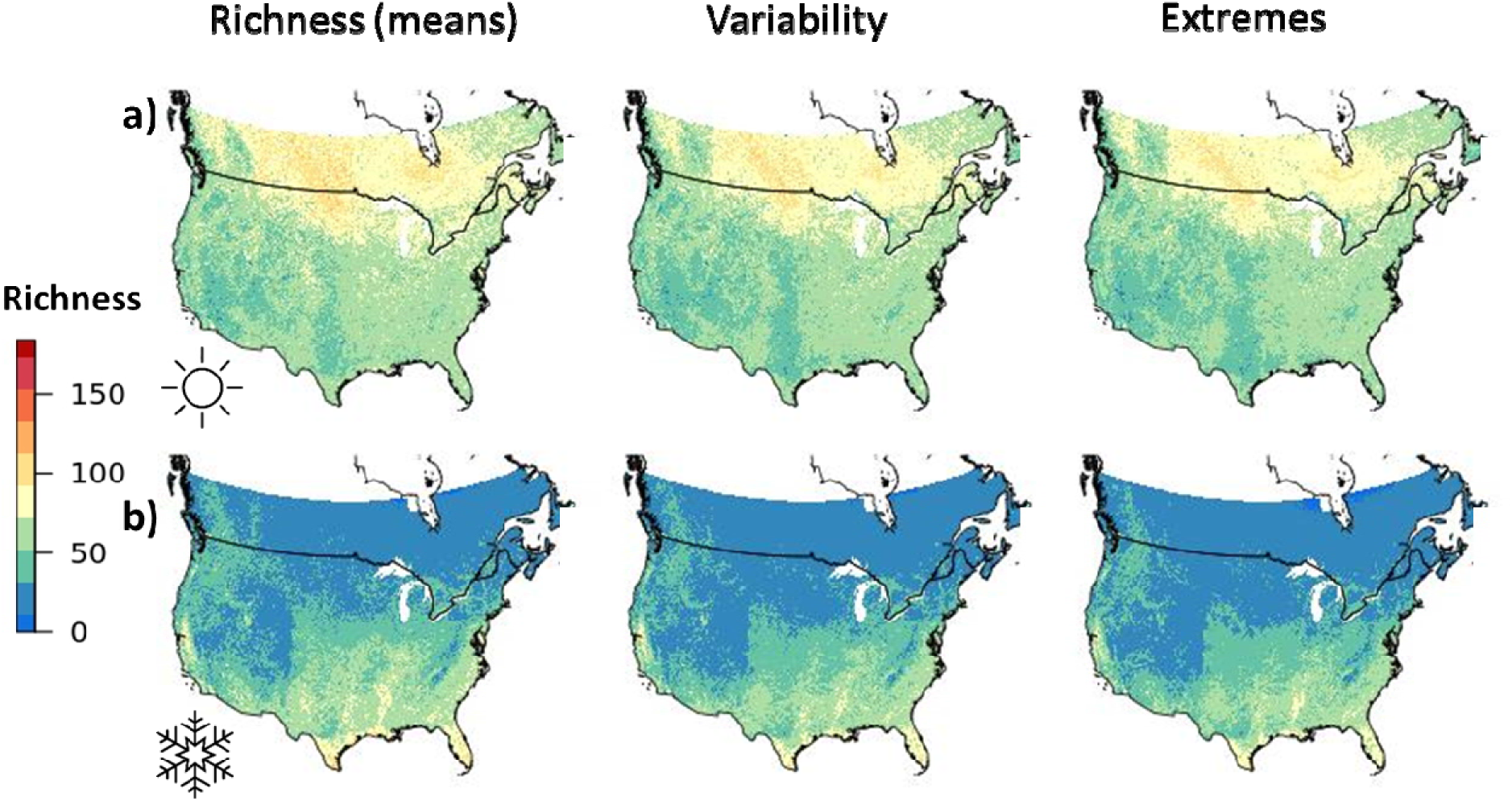
Avian species richness across models containing climate means, variability, and extremes. Predicted richness based on stacked models containing (left) climate means, (middle) variability, and (right) extremes for (top) summer and (bottom) winter.

**Figure S9.**
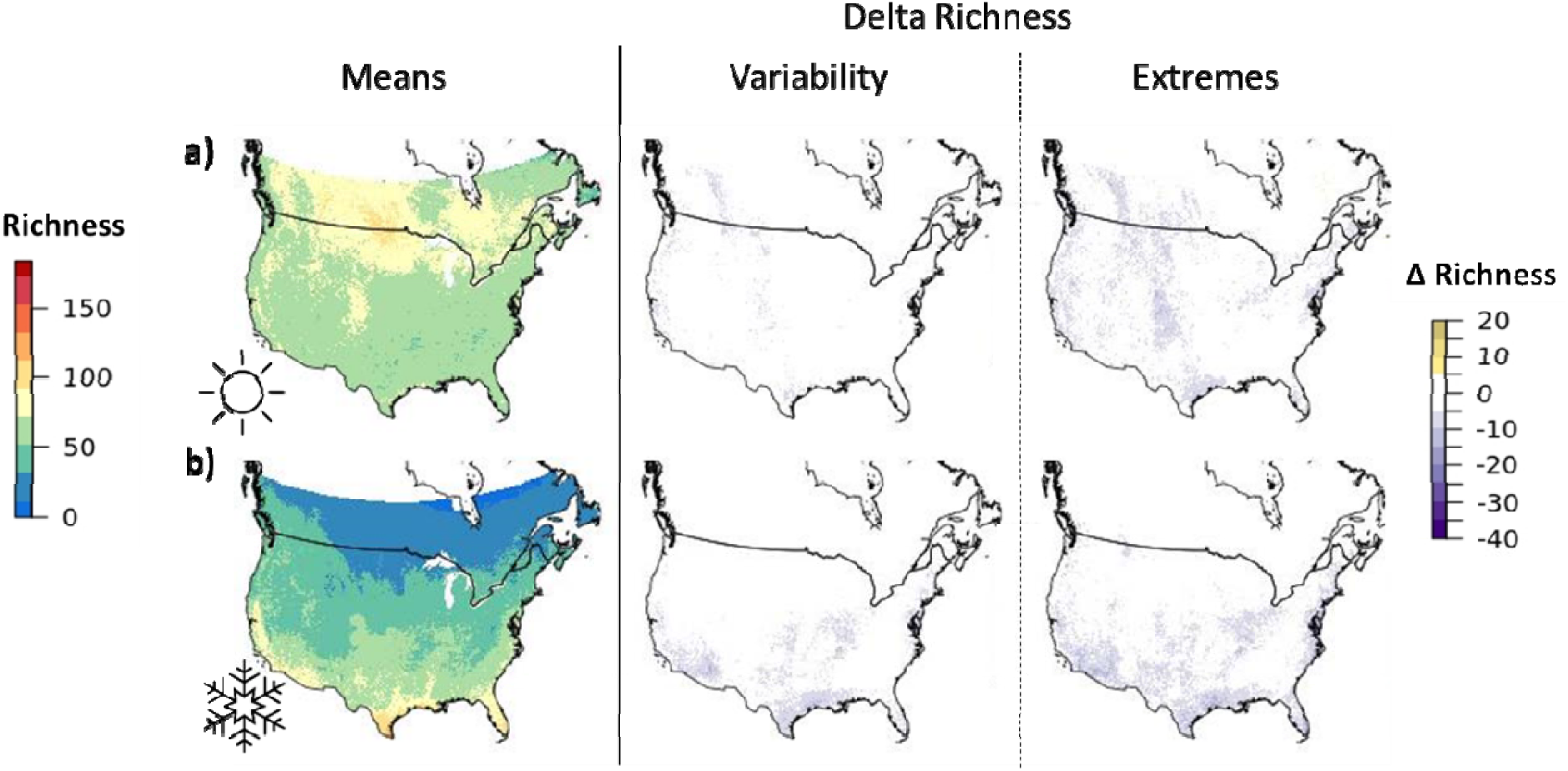
Avian species richness across models containing climate means, variability, and extremes based on relative occurrence rate. Left panels show predicted richness based on stacked relative occurrence rate estimates containing only climate means for (a) summer and (b) winter. Middle and right panels show the difference between predicted richness based on models containing climate variability and extremes compared with means only, with darker purple areas experiencing a larger reduction in estimated richness and yellow areas experiencing an increase in richness.

**Figure S10.**
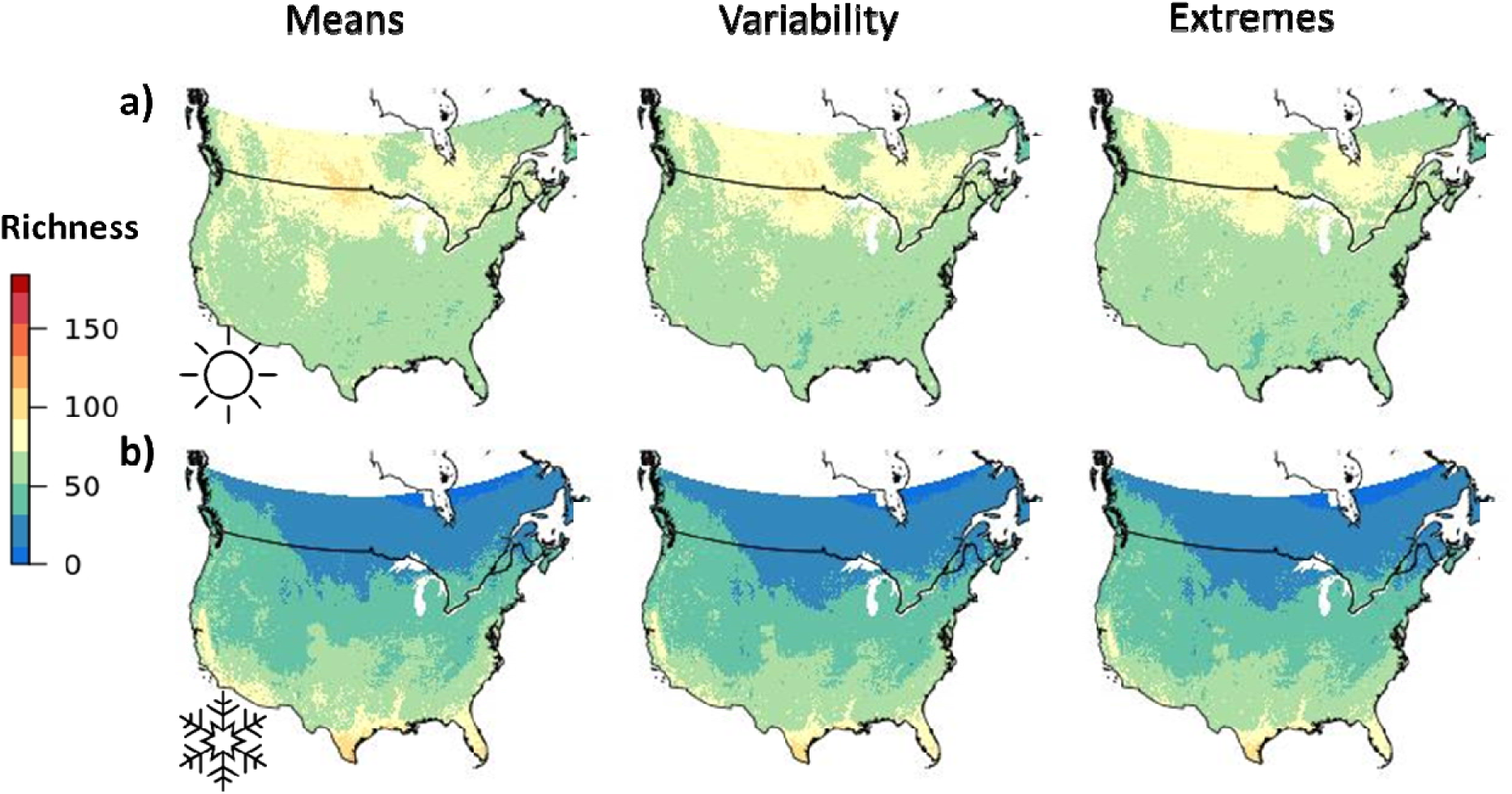
Avian species richness across models containing climate means, variability, and extremes based on relative occurrence rate. Predicted richness based on stacked relative occurrence rate estimates containing (left) climate means, (middle) variability, and (right) extremes for (top) summer and (bottom) winter.

## Supplementary Tables

**Table S1.** Cross-species mean and SD of Moran’s I values to assess spatial autocorrelation in training and testing sets split via spatial-block cross-validation.

| Season | Training mean Moran's I | Training SD | Testing mean Moran's I | Testing SD |
| --- | --- | --- | --- | --- |
| summer | 0.141 | 0.160 | 0.144 | 0.143 |
| winter | 0.127 | 0.127 | 0.139 | 0.145 |

**Table S2.**
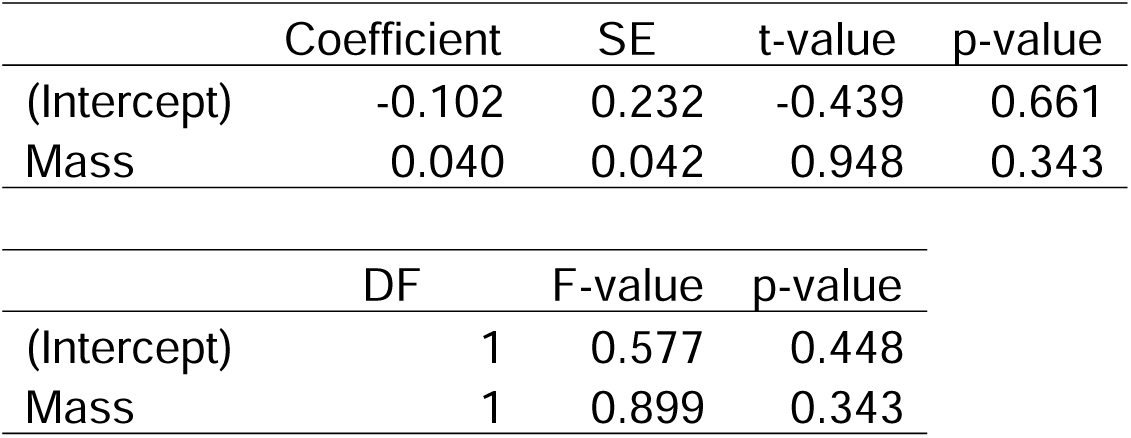
Results of the final phylogenetic least-squares model predicting change in range area between *Means* and *Variability* models across species. Other covariates were dropped during the model selection process. An ANOVA fit to this model is also included.

**Table S3.** Results of the final phylogenetic least-squares model predicting change in proportion of range area predicted beyond expert range boundaries between *Means* and *Variability* models across species. Other covariates were dropped during the model selection process. An ANOVA fit to this model is also included.

|  | Coefficient | SE | t-value | p-value |
| --- | --- | --- | --- | --- |
| (Intercept) | -0.410 | 0.263 | -1.558 | 0.120 |
| Mass | 0.102 | 0.044 | 2.290 | 0.022 |

|  | DF | F-value | p-value |
| --- | --- | --- | --- |
| (Intercept) | 1 | 0.520 | 0.471 |
| Mass | 1 | 5.245 | 0.022 |

**Table S4.** Results of the final phylogenetic least-squares model predicting change in range area between *Means* and *Extremes* models across species. Other covariates were dropped during the model selection process. An ANOVA fit to this model is also included.

|  | Coefficient | SE | t-value | p-value |
| --- | --- | --- | --- | --- |
| (Intercept) | -0.460 | 0.236 | -1.946 | 0.052 |
| Mass | 0.105 | 0.043 | 2.472 | 0.014 |

|  | DF | F-value | p-value |
| --- | --- | --- | --- |
| (Intercept) | 1 | 0.149 | 0.700 |
| Mass | 1 | 6.110 | 0.014 |

**Table S5.** Results of the final phylogenetic least-squares model predicting change in proportion of range area predicted beyond expert range boundaries between *Means* and *Extremes* models across species. Other covariates were dropped during the model selection process. An ANOVA fit to this model is also included.

|  | Coefficient | SE | t-value | p-value |
| --- | --- | --- | --- | --- |
| (Intercept) | -0.473 | 0.212 | -2.235 | 0.026 |
| Mass | 0.104 | 0.038 | 2.755 | 0.006 |

|  | DF | F-value | p-value |
| --- | --- | --- | --- |
| (Intercept) | 1 | 0.072 | 0.789 |
| Mass | 1 | 7.590 | 0.006 |

## Appendices

**Appendix 1.** Modeling information, estimated range size, model performance under cross-validation, and variable importance scores for 481 species in summer and 486 species in winter.

